# The tomato DELLA protein PROCERA promotes ABA responses in guard cells by upregulating ABA transporter

**DOI:** 10.1101/2020.04.22.056010

**Authors:** Hagai Shohat, Natanella Illouz-Eliaz, Yuri Kanno, Mitsunori Seo, David Weiss

## Abstract

Plants reduce transpiration to avoid drought stress by stomatal closure. While abscisic acid (ABA) has a central role in the regulation of stomatal closure under water-deficit conditions, we demonstrated in tomato that the gibberellin (GA) response inhibitor, the DELLA protein PROCERA (PRO), promotes ABA-induced stomatal closure and gene transcription in guard cells. To study how PRO affects stomatal closure, we performed RNAseq analysis of isolated guard cells and identified the ABA transporters *ABA-IMPORTING TRANSPORTER1.1* (*AIT1.1*) and *AIT1.2*, also called in Arabidopsis *NPF4.6*, as upregulated by PRO. Tomato has four *AIT1* genes, but only *AIT1.1* and *AIT1.2* were upregulated by PRO, and only *AIT1.1* exhibited high expression in guard cells. Functional analysis of *AIT1.1* in yeast confirmed its activity as an ABA transporter, possibly importer. CRISPR-Cas9-defrived *ait1.1* mutant exhibited increased transpiration, larger stomatal aperture and reduced response to ABA. Moreover, *ait1.1* suppressed the promoting effects of PRO on ABA-induced stomatal closure and gene expression in guard cells. The negative interaction between GA and ABA has been studied for many years in numerous plant species. These studies suggest that the crosstalk is mediated by changes in hormone biosynthesis and signaling. Our results suggest that it is also mediated by changes in hormone transport.

**One-sentence Summary:** The tomato DELLA protein PROCERA promoted abscisic acid-induced stomatal closure and gene expression by upregulating the expression of the ABA transporter *ABA-IMPORTING TRANSPORTER* 1 in guard cells.

## Introduction

The growth promoting hormone gibberellin (GA) regulates central developmental processes throughout the plant life cycle, from germination to stem elongation, leaf expansion, flowering and fruit development (Yamaguchi, 2008). The output of GA activity on plant development and response to the environment depends on complex interactions with other hormones (Weiss and Ori, 2007). The negative interaction between GA and the stress hormone abscisic acid (ABA) has been studied for many years in numerous plant species. In seeds for example, it regulates dormancy vs. germination; high ABA to GA ratio promotes dormancy, whereas the opposite promotes germination (Razem et al., 2006; Weiss and Ori, 2007).

The nuclear DELLA proteins suppress almost all GA responses by interacting with various transcription factors (Hauvermale et al., 2012; Locascio et al., 2013). When GA binds to its receptor GIBBERELLIN-INSENSITIVE DWARF1 (GID1), it increases the affinity of the latter to DELLA. The generation of GID1-GA-DELLA complex leads to DELLA degradation via the ubiquitin-proteasome pathway, which is mediated by the F-box protein SLEEPY1 (SLY1) (Sasaki et al., 2003; Dill et al., 2004; Griffiths et al., 2006; Harberd et al., 2009; Hauvermale et al., 2012). DELLA destruction in the proteasome leads to transcriptional reprogramming and activation of GA responses. The ability of DELLA to interact with numerous transcription factors is a key factor in the crosstalk between GA and other hormones. For example, DELLA interaction with JASMONATE ZIM DOMAIN (JAZ) proteins mediates the effect of GA on jasmonic acid activity (Hou et al., 2010), and its interaction with BRASSINAZOLE-RESISTANT 1 (BZR1) mediates the crosstalk with brassinosteroids (Li et al., 2012).

The N-terminal region of DELLA (the DELLA domain) is important for the interaction with GID1 and therefore, mutations in this region interfere with the interaction (Harberd et al., 2009). These dominant, gain-of-function mutations stabilize DELLA, leading to constitutive inhibition of GA responses. The C-terminal region of DELLA (the GRAS domain) plays a major role in repressing GA responses by interacting with numerous transcription factors (Yoshida et al., 2014). Mutations in the C-terminal region are recessive, and exhibit constitutive GA responses (Sun and Gubler, 2004; Harberd et al., 2009). Tomato (*Solanum lycopersicum*) has one DELLA protein, called PROCERA (PRO) (Jasinski et al., 2008; Livne et al., 2015). The tomato loss-of-function mutants *pro* and *pro*^*ΔGRAS*^ are tall and exhibit increased GA responses (Van Tuinen et al., 1999; Bassel et al., 2008; Fleishon et al., 2011; Livne et al., 2015), whereas the gain-of-function *proGF* mutant is dwarf due to constitutive inhibition of GA responses (Zhu et al., 2019).

The crosstalk between GA and ABA has been studied for many years in numerous plant species (Piskurewicz et al., 2008; Liu et al., 2018, Shu et al., 2018). The transcription factor ABSCISIC ACID-INSENSITIVE 4 (ABI4) promotes seed dormancy in Arabidopsis by the suppression of GA accumulation and the promotion of ABA biosynthesis (Shu et al., 2013). In *Sorghum, SbABI4* promotes the transcription of the GA deactivating gene *SbGA2ox3* (Cantoro et al., 2013). Moreover, the transcription factor GERMINATION INSENSITIVE TO ABA MUTANT 2 (GIM2) promotes GA biosynthesis while reducing ABA production (Xiong et al., 2017). In Arabidopsis seeds, DELLA promotes the expression of the RING ubiquitin E3 ligase XERICO that is involved in ABA accumulation. It also increases the expression of the transcription factor *ABI5* that inhibits seed germination, and interacts with the ABA signaling components ABI3 (Lim et al., 2013). ABA, in turn, stabilizes the Arabidopsis DELLA protein REPRESSOR OF GA1-3 LIKE-2 (RGL2) and inhibits GA signaling (Piskurewicz et al., 2008). In tomato, the lack of DELLA activity in seeds suppresses desiccation tolerance due to inhibition of ABA-induced gene expression (Livne et al., 2015). Taken together, these studies suggest that GA and ABA negatively interact at the hormone biosynthesis and signaling levels (Shu et al., 2018).

Previously we suggested a crosstalk between GA/DELLA and ABA in tomato guard cells (Nir et al., 2017). Transgenic tomato plants overexpressing the Arabidopsis or the tomato stable DELLA mutant proteins *rgaΔ17* or *proΔ17*, respectively, exhibited reduced stomatal aperture and transpiration. The effects of *pro*Δ*17* on stomatal closure and water loss were suppressed in the ABA-deficient *sitiens* (*sit*) mutant, indicating that these effects of DELLA are ABA-dependent (Nir et al., 2017). We found that DELLA promotes ABA responses, including ABA-induced stomatal closure and reactive oxygen species (ROS) accumulation following ABA application to guard cells. Since DELLA is a transcription regulator, it is yet unclear how PRO affects ABA-induced stomatal closure. Since PRO did not affect ABA accumulation, we speculated that it affects ABA signaling or uptake into guard cells via transcriptional regulation of ABA signaling component or transporter genes (Nir et al., 2017).

Several ABA transporters were identified and characterized in Arabidopsis, including the ATP-BINDING CASSETTE (ABC) transporters ABCG25, ABCG40 and ABA-IMPORTING TRANSPORTER 1 (AIT1), also called NITRATE TRANSPORTER 1.2 (NRT1.2) or NRT1/PTR TRANSPORTER FAMILY 4.6 (NPF4.6). ABCG25 is expressed in vascular tissues and functions as an ABA exporter (Kuromori et al., 2010). ABCG40, is an ABA importer that was localized to the guard-cell plasma membrane (Kang et al., 2010). *AIT1* transcripts were found in the vascular tissues of inflorescence stems and the *ait1* mutant exhibited increased water loss due to open stomata (Kanno et al., 2012). Loss of ABCG25 increases the sensitivity to ABA whereas the loss of ABCG40 and AIT1 reduced the sensitivity to the hormone (Kuromori et al., 2018).

Here we studied the mechanism by which the tomato DELLA protein PRO increases ABA responses in guard cells. RNAseq analysis of isolated guard cells identified the ABA transporter AIT1.1 as upregulated by PRO. The loss of AIT1.1 suppressed the effect of PRO on guard-cell ABA responses.

## Results

### PRO promotes ABA responses in guard cells

To support our previous suggestion that DELLA promotes ABA responses in guard cells (Nir et al., 2017), we tested the effect of PRO on ABA-inhibition of transpiration and ABA-induced gene expression in guard cells. Thermal imaging of M82 and transgenic plants overexpressing the stable DELLA protein *pro*Δ*17* (*35S:proΔ17*, Nir et al., 2017) showed higher leaf-surface temperature in the transgenic line following the application of ABA, indicating lower transpiration rate (Figure 1A). To examine the effect of DELLA on ABA-induced transcription, we have generated transgenic M82 plants expressing the reporter *β-glucuronidase* (*GUS*) gene under the regulation of the Arabidopsis ABA-induced promoter *MAPKKK18* (Okamoto et al., 2013). The transgene was then introgressed into *35S:proΔ17* plants by crosses. The GUS signal in ABA-treated leaves was observed in guard cells and was stronger in *35S:proΔ17* compared to M82 (Figure 1B). These results suggest that PRO promotes ABA physiological and transcriptional responses in guard cells.

**Figure 1.**
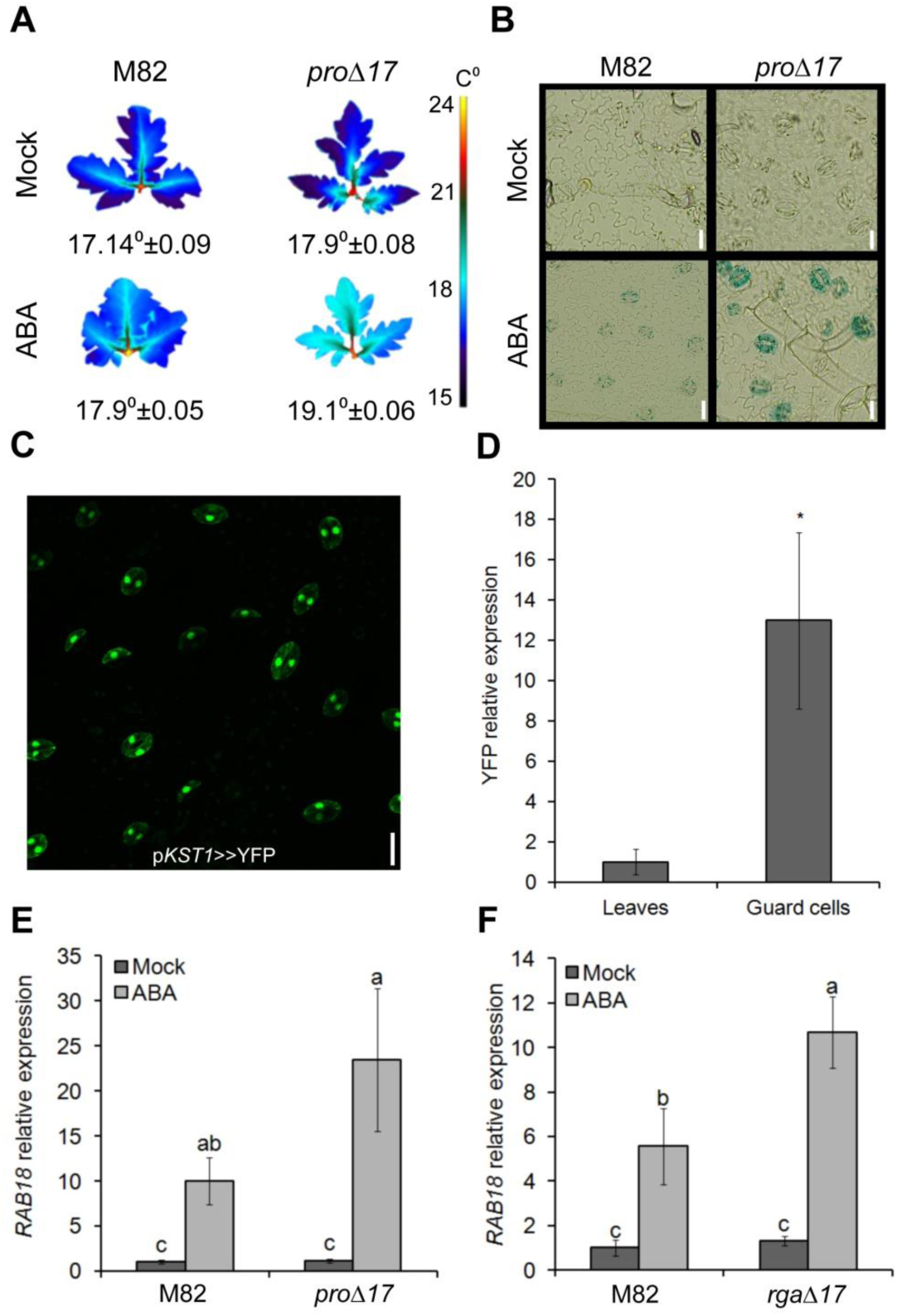
PRO promotes ABA responses in guard cells. **A.** Thermal imaging of leaves (third leaf) taken M82 and *35S:proΔ17* treated or not (Mock) with 10 μM ABA. Number below leaves are the calculated leaf-surface temperature and the values are means of three plants, measured 20 times ± SE. **B.** Representative images of GUS staining of epidermal peels treated or not (Mock) with 10 μM ABA. Peels were taken from the fourth leaves of M82 and *35S:proΔ17* expressing the reporter GUS under the regulation of the *MAPKKK18* promoter (Scale Bar = 20µm). **C**. YFP signal in guard cells of *pKST1>>YFP* trans-activated epidermal peel (Scale Bar = 20µm). **D**. YFP expression in whole leaf tissue and guard cell enriched samples. Values are means of four biological replicates ± SE. Stars above the columns represents significant differences between respective treatments (Student’s t test, P<0.05). **E and F.** qRT-PCR analysis of *RAB18* expression in guard cell enriched samples taken from M82 and *35S:proΔ17* (E) or *35S:rgaΔ17* (F). Values in E and F are means of four biological replicates ± SE. Lowercase letters represent significant differences between lines and treatments (Student’s t-test, P<0.05). Small letters above the columns represents significant differences between respective treatments (Tukey–Kramer HSD, P<0.05). The value for leaves in D was set to 1 and the value for M82 Mock in E and F was set to 1.

Since DELLA is a transcription regulator, we hypothesized that PRO affects transpiration and stomatal movement by regulating the expression of ABA/stomatal-related genes in guard cells. To study the interaction between DELLA and ABA in the regulation of gene expression, we first developed a rapid and efficient guard-cell isolation protocol to minimize the effect of the isolation on gene expression (see Material and methods). To validate the procedure, we have used plants expressing the YELLOW FLUORESCENT PROTEIN (YFP) under the guard-cell-specific promoter *KST1* (Figure 1C; Nir et al., 2017). qRT-PCR analysis of RNA extracted from whole leaf tissue or isolated guard cells, showed ca. 13-fold higher YFP expression in the guard-cells enriched samples (Figure 1D). We then used this procedure to test the response of the ABA-induced gene *RAB18* (Nir et al., 2017) to ABA in guard-cell enriched samples taken from M82, *35S:pro*Δ*17* and *35S:rga*Δ*17*. The expression of *RAB18* following ABA treatment was higher in the transgenic lines (Figure 1E and F).

### Global expression response to PRO in guard cells

We next explored the mechanism by which DELLA promotes ABA-induced stomatal closure. To this end, we examined the global effect of PRO on guard cells transcriptional activity, by performing RNAseq analysis to guard-cells enriched samples taken from M82, *35S:pro*Δ*17* and *pro*. Using a 2-fold increase or decrease cutoff (adjusted P value for multiple comparisons ≤0.05) we identified 162 PRO-regulated genes (81 upregulated and 81 downregulated genes, Figure 2A, Supplemental Data set 1, https://www.ncbi.nlm.nih.gov/geo/query/acc.cgi?acc=GSE143999). We then searched for differentially expressed genes related to stomatal closure and/or ABA. Among these genes (Supplemental Table 1), we identified three putative ABA transporters; the tomato homologs of the Arabidopsis *ABCG40* and two *AIT1* genes, also called in Arabidopsis *NPF4.6* or *NRT1.2* (Supplemental Figure 1; Kanno et al., 2012). We named the tomato proteins AIT1.1 and AIT1.2. The expression of *ABCG40* and *AIT1.2* in the RNAseq analysis was extremely low (Supplemental Table 1). It is worth noting that although we reported previously that two ABA receptors *PYRABACTIN RESISTANCE 1(PYR1*) and *PYR1-like8-1* (*PYL8-1*) are upregulated in *35S:pro*Δ*17* (Nir et al., 2017), in the RNAseq analysis we did not find them among the differentially expressed genes.

**Figure 2.**
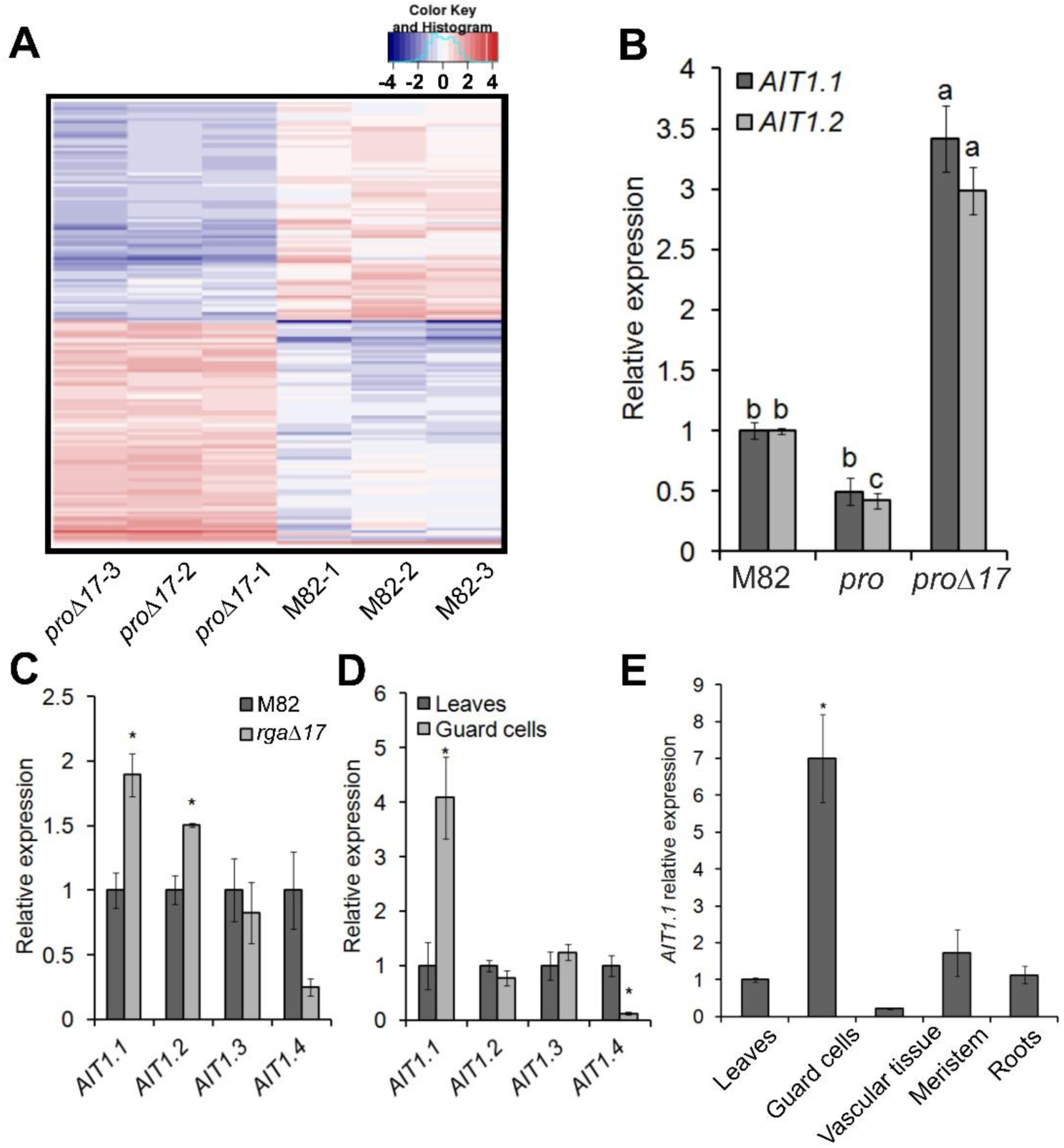
RNAseq analysis identified the ABA transporter AIT1.1 as upregulated by PRO in guard cells. **A.** Clustered heatmap of PRO regulated genes (*proΔ17* vs. M82, three samples each) generated from RNAseq and shows 81 PRO upregulated and 81 downregulated genes. Genes were grouped based on their pattern of expression. Coloring of the genes is according to the color bar on the upper right side (Log2 fold change). The complete list of PRO-regulated genes are provided in Supplemental dataset 1. **B.** qRT-PCR analysis of *AIT1.1* and *AIT1.2* expression in M82, *pro* and *35S*:*pro*Δ*17* (*proΔ17*) isolated guard cell. Values are means of three biological replicates ± SE. Lowercase letters represent significant differences between lines (Tukey–Kramer HSD, P < 0.05). **C.** Expression of all tomato *AIT1* genes in M82 and *35S*:*rgaΔ17* (*rgaΔ17*) isolated guard cell. **D.** Expression of all tomato *AIT1* genes in leaves and isolated guard cells. **E.** Expression of AIT1.1 in different tissues. Values in C, D and E are means of four replicates ± SE. Stars above the columns represents significant differences between respective treatments (Student’s t test, P<0.05). The values for M82 in B and C was set to 1 and the values for leaves in D and E was set to 1.

We first validated the results of the RNAseq for the effect of PRO on the expression of *ABCG40, AIT1.1* and *AIT1.2* in guard-cell enriched samples by qRT-PCR. This analysis did not confirm the effect of PRO on *ABCG40* (Supplemental Figure 2), but it confirmed its effect on *AIT1.1* and *AIT1.2*. These genes were up- and downregulated in *35S:pro*Δ*17* and *pro*, respectively (Figure 2B). Tomato has 4 *AIT1* homologs that we named *AIT1.1* to *AIT1.4* (Supplemental Figure 1). We analyzed the expression of all four *AIT1* homologs in M82 and *35S:rga*Δ*17* guard cells and only *AIT1.1* and *AIT1.2* were upregulated by DELLA in guard cells (Figure 2C). We then analyzed the expression of all *AIT1s* in M82 guard-cell enriched samples compared to whole leaf tissue, and only *AIT1.1* exhibited significantly higher expression in guard cells (Figure 2D). Kanno et al. (2012), found that the Arabidopsis *AIT1* is expressed in the vascular tissue, using reporter line. We used the same approach and generated transgenic M82 plants expressing the GUS reporter under the regulation of *AIT1.1* promoter (∼1400 bp upstream the start codon). The tomato *AIT1*.*1* also showed high GUS activity in vascular tissues (Supplemental Figure 3), but not in guard cells. We therefore analyzed *AIT1.1* expression in various tissues of M82 plants by qRT-PCR (mature leaves, shoot apices, vascular tissue, guard cells and roots) and found the highest expression in guard cells (Figure 2E). These results suggest that the promoter used to express GUS did not provide the authentic spatial expression pattern.

### PRO promotes ABA responses via the ABA transporter *AIT1.1*

Functional analysis of the Arabidopsis AIT1 protein in yeast cells, suggests that it operates as an ABA importer (Kanno et al., 2012). When expressed in yeasts, the tomato AIT1.1 induced the interactions between the Arabidopsis ABA receptor PYR1 and the protein phosphatase 2C (PP2C) ABI1 under a relatively low ABA concentration (0.5 µM) in the growth media, as the Arabidopsis AIT1 protein did, while the interactions were not observed in the absence of ABA (Figure 3A). We further confirmed that AIT1.1 mediated cellular ABA uptake by directly quantifying the molecules taken into the yeast cells by liquid chromatography mass-spectrometry (LC-MS/MS) (Figure 3B). We also examined the substrate selectivity of AIT1.1 against several other hormones. It appeared that AIT1.1 transported GA1, GA4 and indole-3-acetic acid (IAA) to some extent, however, the selectivity was much lower compared to ABA (Figure 3B).

**Figure 3.**
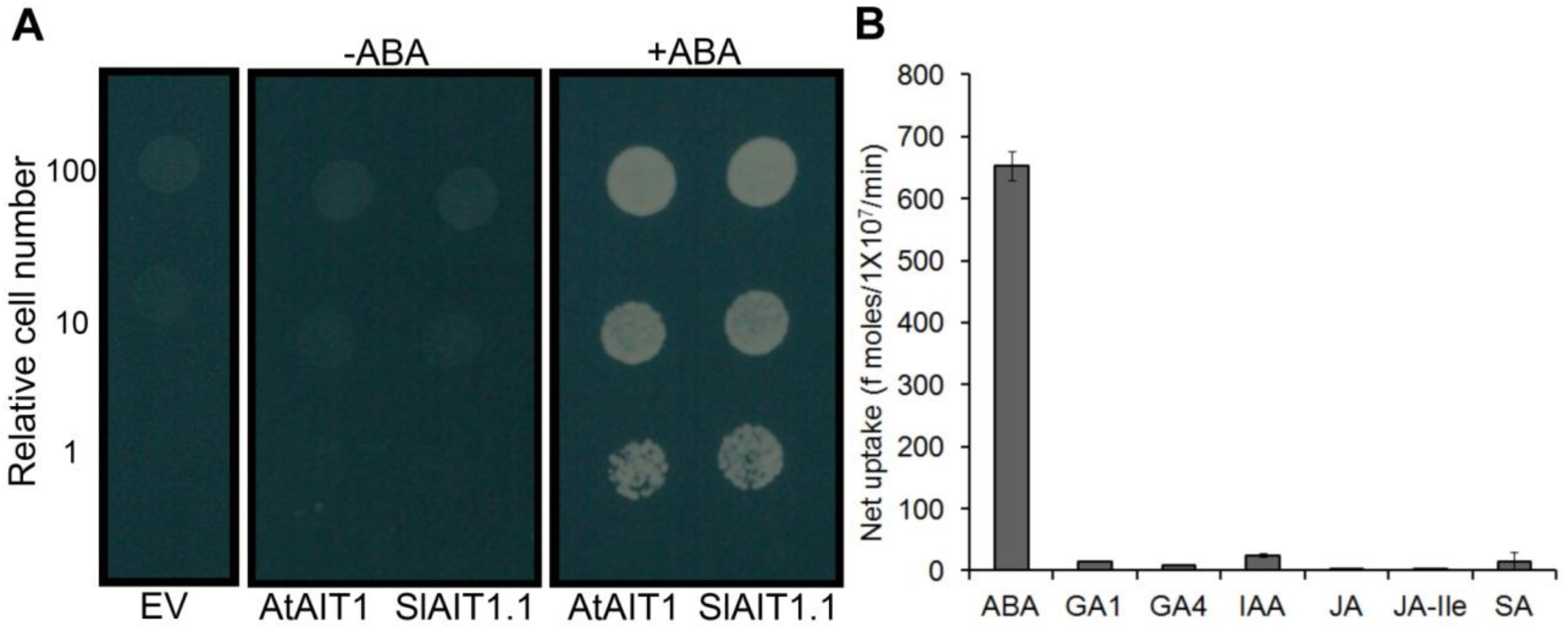
AIT1.1 mediates ABA uptake into yeast cells. **A.** Effects of AIT1.1 on the interactions between the ABA receptor and PP2C. Tomato AIT1.1 (*SlAIT1.1*) or Arabidopsis AIT1 (*AtAIT1*) was expressed in yeast containing a yeast two-hybrid system with the Arabidopsis PYR1 ABA receptor fused to the GAL4 DNA binding domain and the ABI1 protein phosphatase fused to the GAL4 activation domain, and the cells were inoculated on selection media (SD, -Leu, -Trip, -Ura, -His) containing 0.5 µM ABA (+ABA) or without ABA (-ABA). An empty vector (EV) was transformed as a negative control. Photos were taken 3 days after inoculation. **B.** Hormone transport activities of AIT1.1. Yeast cells expressing tomato AIT1.1 were incubated with solutions containing 10 µM ABA, GA_1_, GA_4_, indole-3-acetic acid (IAA), jasmonic acid (JA), jasmonoyl-isoleucine (JA-Ile) or salicylic acid (SA), and the amounts of compounds taken into the cells were quantified with LC-MS/MS.

Although *AIT1.2* was upregulated by PRO, it showed very low expression levels (ca. 20 fold lower than *AIT1.1*, Supplemental Table 1) and its expression in guard cells was similar to whole leaf tissue. We therefore focused further on *AIT1.1*. We generated CRISPR-Cas9 derived *ait1.1* mutant and obtained two independent alleles. Both mutations caused a frame shift and premature stop codon (Supplemental Figure 4). Homozygous plants of the two lines exhibited mild growth suppression (Supplemental Figure 5A and B). Thermal imaging showed lower leaf-surface temperature in *ait1.1*, suggesting that transpiration in *ait1.1* is higher than in M82 (Figure 4A). *ait1.1* exhibited larger stomatal aperture (Figure 4B and Supplemental Figure 5C) and partial inhibition of stomatal closure in response to application of 10μM ABA to epidermal peels (Figure 4C).

**Figure 4.**
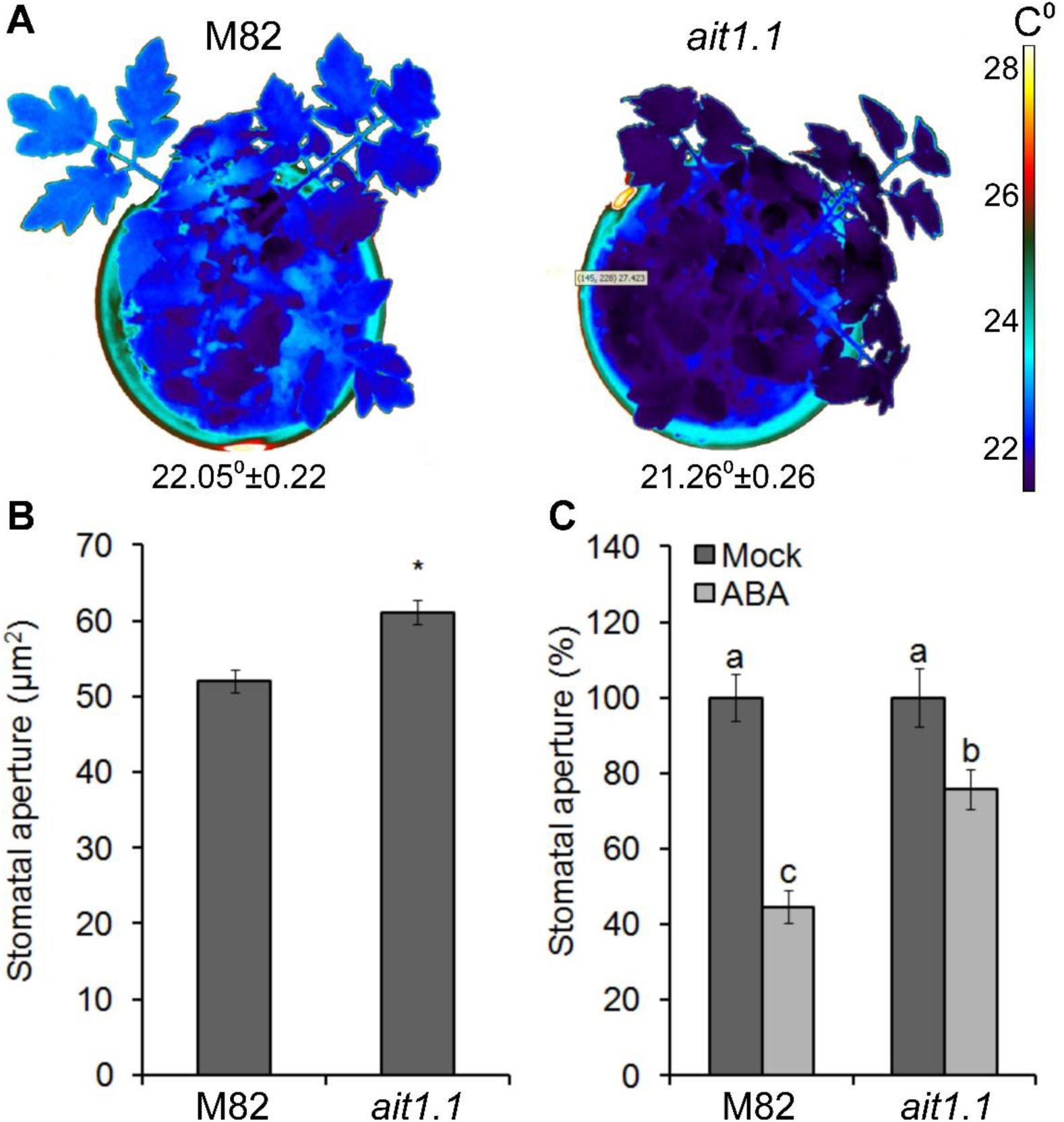
Loss of the ABA transporter *AIT1.1* increased transpiration and inhibited ABA responses in guard cells. **A.** Termal imaging of M82 and CRISPR-Cas9 derived *ait1.1* mutant. Numbers below plants are the leaf-surface temperature and the values are means of three replicated each measured 20 times ± SE. **B**. Stomatal aperture measured on imprints of abaxial epidermis taken at 11:00 h. Values are means of four replicates, each with ca. 100 measurements (stomata) ± SE. Stars above the columns represents significant differences between respective treatments (Student’s t test, P<0.05).**C**. Stomatal aperture in M82 and *ait1.1* epidermal peels treated or not treated (Mock) with 10 μM ABA. One hour after the ABA treatment stomatal aperture was measured. Values are mean precentage of mock of four replicates, each with ca. 100 measurements (stomata) ± SE. Different letters above the columns represent significant differences between lines and treatments (Tukey–Kramer HSD, P < 0.05).

We then introgressed the *35S:rga*Δ*17* transgene into *ait1.1* background by crosses and generated homozygous *ait1.1* plants overexpressing 35S:*rga*Δ*17*. We confirmed the presence of the transgene (*35S:rga*Δ*17*) by the phenotype (shorter stem and smaller, darker and more serrated leaves, Supplemental Figure 6), and the *ait1.1* mutation by sequencing. The loss of *AIT1.1* suppressed the effect of stable DELLA overexpression on transpiration as indicated by the lower leaf-surface temperature in *35S:rgaΔ17 ait1.1* plants compared to *35S:rga*Δ*17* (Figure 5A and B). In addition, stomatal response to ABA in epidermal peels, and the expression of the ABA-induced gene *RAB18* were suppressed in *35S:rgaΔ17 ait1.1* compared to *35S:rga*Δ*17* (Figure 5C and D). These results suggest that AIT1.1 is required for DELLA to promote ABA responses in guard cells.

**Figure 5.**
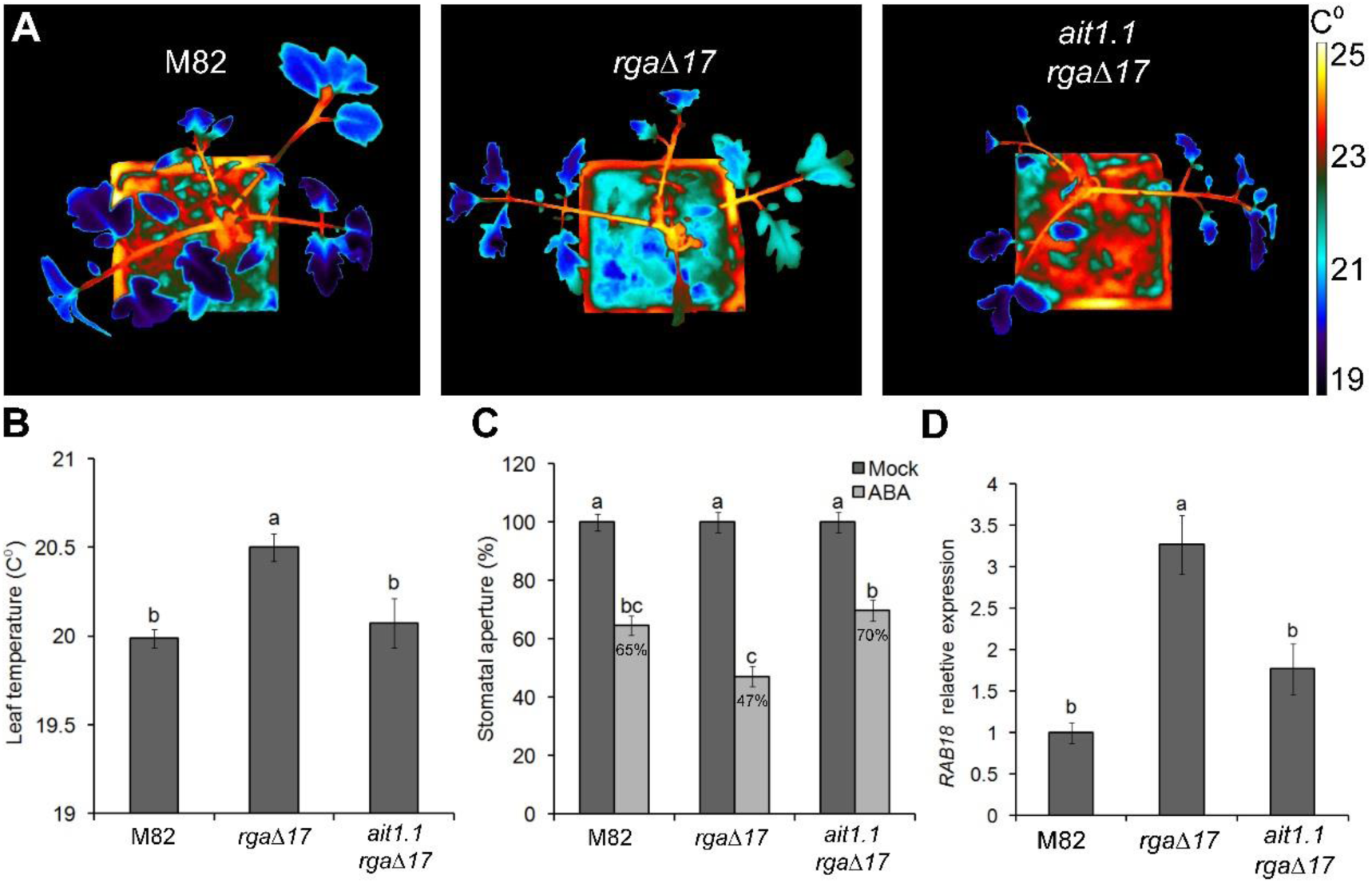
*ait1.1* suppressed the effect of PRO on ABA responses in guard cells. **A.** Termal imaging of M82, *35S:rgaΔ17* (*rgaΔ17*) and *rgaΔ17 ait1.1*. **B**. Leaf-surface temperature of M82, *rgaΔ17* and *rgaΔ17 ait1.1*. plants. Values are means of three replicated measured 20 times ± SE. Different letters above the columns represent significant differences between lines (Student’s t test, P<0.05). **C.** Stomatal aperture in M82, *rgaΔ17* and *rgaΔ17 ait1.1* epidermal peels treated or not treated (Mock) with 10 μM ABA. One hour after the ABA treatment stomatal aperture was measured. Values are mean precentage of mock of four replicates, each with ca. 100 measurements (stomata) ± SE. Different letters above the columns represent significant differences between lines and treatments (Tukey–Kramer HSD, P < 0.05). **D.** qRT-PCR analysis of *RAB18* expression in M82, *rgaΔ17* and *rgaΔ17 ait1.1* isolated guard cells. Values are means of four biological replicates ± SE. Different letters above the columns represent significant differences between lines (Student’s t test, P<0.05). The values for M82 was set to 1.

## Discussion

Drought avoidance is a major plant-adaptation strategy to survive transient water-deficit conditions (Kooyers, 2015). To avoid drought stress, plants close their stomata to reduce transpiration and can use the available water in the soil more slowly and for longer time before the next rain is coming. While ABA has a major role in the regulation of transpiration under water-deficit stress, accumulating evidence suggest that GA antagonizes these ABA responses. Increased GA levels promotes stomatal opening in *Commelina benghalensis* and *Vicia faba* (Santakumari and Fletcher, 1987; Göring et al., 1990). Reduced GA activity and DELLA accumulation suppress canopy expansion and xylem hydraulic conductivity and promotes stomatal closure, all leading to lower transpiration (Nir et al., 2014; Nir et al., 2017; Illouz-Eliaz et al., 2020). It was suggested that water deficiency inhibits GA accumulation to promote adaptation to drought (Colebrook et al., 2014).

Since ABA-induced stomatal closure is mediated by the phosphorylation of ion channels and not by activation of gene transcription (Cutler et al., 2010; Kim et al., 2010; Cai et al., 2017), it was unclear how the transcriptional regulator PRO affects ABA-induced stomatal closure. We hypothesized that PRO affects the transcription of either ABA signaling component or ABA transporter (Nir et al., 2017). RNAseq analysis of isolated guard cells (M82, *35S:pro*Δ*17* and *pro*) identified the ABA transporter *AIT1.1* as upregulated by PRO. The Arabidopsis *AIT1*, also called *NRT1.2* or *NPF4.6*, is an ABA importer (Kanno et al., 2012). This gene is also upregulated by DELLA in the Arabidopsis shoot apical meristems (Serrano-Mislata et al., 2017). The Arabidopsis *ait1* mutant exhibited increased water loss due to open stomata (Kanno et al., 2012), however, expression analysis (based on reporter line), suggests that AIT1 is active in the vascular tissue of inflorescence stems, but not in stomata. The high expression levels of the tomato *AIT1.1* in guard cells and the open stomata in the *ait1.1* mutant, suggest that in tomato this protein function as an ABA transporter, possibly as an importer, in guard cells. It is rather clear that AIT1.1 is not the only ABA transporter in tomato guard cells. However, among the AIT1 group of transporters, AIT1.1 is the most dominant one, based on its expression level. In Arabidopsis, an importer from another group, the ABC transporter ABCG40 is active in guard cells (Kang et al., 2010). We identified the *ABCG40* homolog in the RNAseq of isolated guard cells as PRO-upregulated. However, this was not confirmed by qRT-PCR analysis. Thus, ABCG40 may contribute to ABA uptake into tomato guard cells, but probably does not mediate the effect of PRO on guard-cell ABA responses.

The up- and down-regulation of *AIT1.1* by stable PRO and *pro* loss-of-function, respectively, and the suppression of PRO-promoted ABA responses in *ait1.1* guard cells, suggest that *AIT1.1* mediates the effect of PRO on ABA-induced stomatal closure. It is possible that water deficiency reduces the levels of active GAs in guard cells, leading to PRO accumulation. PRO promotes the expression of the ABA importer *AIT1.1*, facilitating ABA uptake into guard cells, leading to faster stomatal closure.

The crosstalk between GA and ABA has been investigated for many years (Weiss and Ori, 2007; Shu et al., 2018). This crosstalk is largely depends on DELLA; DELLA promotes ABA synthesis and signaling and ABA promotes DELLA stability and therefore, inhibits GA signaling (Achard et al., 2006; Piskurewicz et al., 2008; Lim et al., 2013; Liu et al., 2018). Here we bring evidence that PRO (DELLA) promotes ABA responses in guard cells via the upregulation of the ABA transporter *AIT1.1*, suggesting that the crosstalk between GA and ABA is mediated at multiple levels; hormone biosynthesis, signaling and transport.

## Material and methods

### Plant materials, growth conditions and hormone treatments

Tomato (*Solanum lycopersicum*) plants in M82 background (*sp/sp*) were used throughout this study. The *pro* mutant was in M82 background (Fleishon et al., 2011). The CRISPER-derived *ait1.1* mutant and the transgenic lines 35S:*rgaΔ17* (Livne et al., 2015), 35S:*proΔ17, pKST1*:LhG4 (Nir et al., 2017) and *pMAPKKK18*:GUS were generated in M82 background. Plants were grown in a growth room set to a photoperiod of 12/12-h night/day, light intensity of 200 μmol m^-2^ s^-1^ and 25°C. In other experiments, plants were grown in a greenhouse under natural day-length conditions, light intensity of 700 to 1000 µmol m^-2^ s^-1^ and 18-30°C. The seeds were harvested from ripe fruits and treated with 1% Sodium hypochlorite followed by 1% Na_3_PO_4_ 12H_2_O, and incubated with 10% Sucrose over-night in 37°C. Seeds were stored dry at room temperature. (±)-ABA dissolved in DMSO (Sigma-Aldrich, St. Louis, USA) was applied to plants by spraying.

### CRISPR/Cas9 mutagenesis, tomato transformation and selection of mutant alleles

Four single-guide RNAs (sgRNAs, Supplemental Table 2) were designed to target *AIT1.1* gene, using the CRISPR-P tool (http://cbi.hzau.edu.cn/crispr). Vectors were assembled using the Golden Gate cloning system, as described by Weber et al. (2011). Final binary vectors, pAGM4723, were introduced into *A. tumefaciens* strain GV3101 by electroporation. The constructs were transferred into M82 cotyledons using transformation and regeneration methods described by McCormick (1991). Kanamycin-resistant T0 plants were grown and independent transgenic lines were selected and self-pollinated to generate homozygous transgenic lines. The genomic DNA of each plant was extracted, and genotyped by PCR for the presence of the Cas9 construct. The CRISPR/Cas9-positive lines were further genotyped for mutations using a forward primer to the upstream sequence of the sgRNA1 target and a reverse primer to the downstream of the sgRNA2 target sequence. The target genes in all mutant lines were sequenced. Several homozygous and heterozygous lines were identified and independent mutant lines for each gene were selected for further analysis. The Cas9 construct was segregated out by crosses to M82.

### Molecular cloning/construct and transactivation

To generate the ABA-reporter transgenic plants, the plasmid containing the *AtMAPKKK18* promoter were kindly supplied by Assaf Mosquna (Okamoto et al., 2013). To generate *pMAPKKK18:GUS* the *MAPKKK18* promoter was inserted into the pRITA vector downstream to the GUS start codon into the KpnI and BamHI sites. This construct was then cloned into the pART27 binary vector and introduced to *A. tumefaciens* strain GV3101 by electroporation to generate transgenic M82 tomato plants (as describe above). To specifically express YFP in guard cells, the LhG4 transactivation system (Moore et al., 1998) with the *KST1* promoter was used. *pKST1*:LhG4 used as a driver line and OP:YFP as the responder line. The cross between these lines generated the trans-activated line *pKST1*>>YFP.

### *pAIT1.1:GUS* molecular cloning

The 1380 bp upstream to *AIT1.1* start codon was amplified by PCR (−49 to - 1429). *pAIT1.1* was inserted into the KpnI and BamHI sites in pRITA vector downstream to the GUS start. This construct was then cloned into the pART27 binary vector and introduced to *A. tumefaciens* strain GV3101 by electroporation to generate transgenic M82 tomato plants (as describe above).

### Library constructions and sequencing

Total RNA was extracted from isolated guard cells using RNeasy Plant Mini Kit (Qiagen, Hilden, Germany). Libraries were prepared at the Crown Institute for Genomics (G-INCPM, Weizmann Institute of Science). 500ng of total RNA for each sample was processed using the Inhouse poly A based RNA seq protocol. Libraries were evaluated by Qubit and TapeStation. Sequencing libraries were constructed with barcodes to allow multiplexing of 9 samples on one lane of Illumina HiSeq 2500 machine, using the Single-Read 60 protocol (v4), yielding a median of 29.4M reads per sample (Illumina; single read sequencing).

### Sequence data analysis

Stretches of Poly-A/T and Illumina adapters were trimmed from the reads using cutadapt (Martin et al., 2011) resulting reads shorter than 30bp were discarded. Reads were mapped to the *Solanum lycopersicum* reference genome; release SL3.00 using STAR (Dobin et al., 2013) (with End To End option and out Filter Mismatch Nover Lmax set to 0.04). Annotation file was downloaded from SolGenomics, ITAG release 3.2. Expression levels for each gene were quantified using htseq-count (Andres et al., 2014), using the gtf above (using union mode). Differentially expression analysis was performed using DESeq2 (Love et al., 2014) with the betaPrior, cooksCutoff and independent Filtering parameters set to False. Raw P values were adjusted for multiple testing using the procedure of Benjamini and Hochberg (Bejamini and Hochberg, 1995). The RNA-Seq data discussed in this publication have been deposited in NCBI’s Gene Expression Omnibus (Edgar et al., 2002) and are accessible through GEO Series accession number GSE143999 (https://www.ncbi.nlm.nih.gov/geo/query/acc.cgi?acc=GSE143999).

### RNA extraction and cDNA synthesis

Total RNA extracted by RNeasy Plant Mini Kit (Qiagen). For synthesis of cDNA SuperScript II reverse transcriptase (18064014; Invitrogen, Waltham, MA, USA) and 3 mg of total RNA were used, according to the manufacturer’s instructions.

### qRT-PCR analysis

qRT-PCR analysis was performed using an Absolute Blue qPCR SYBR Green ROX Mix (AB-4162/B) kit (Thermo Fisher Scientific, Waltham, MA, USA). Reactions were performed using a Rotor-Gene 6000 cycler (Corbett Research, Sydney, Australia). A standard curve was obtained using dilutions of the cDNA sample. The expression was quantified using Corbett Research Rotor-Gene software. Three independent technical repeats were performed for each sample. Relative expression was calculated by dividing the expression level of the examined gene by that of *SlACTIN*. Gene to *ACTIN* ratio was then averaged. All primers sequences are presented in Supplemental Table 3.

### Isolation of guard cells

Fully expanded leaves without the central veins were ground twice in a blender containing 100 ml cold distilled water, for 60 sec each time. The blended mixture was poured onto a 100 µm nylon mesh (Sefar, Heiden, Switzerland) and the remaining epidermal peels were rinsed thoroughly with 0.5L of cold deionized water. The peels were then transferred into a 2 ml tubes and froze in liquid nitrogen.

### Microscopy and Confocal imaging analysis

Imaging was done using a confocal laser scanning microscope model SP8 (Leica Microsystem, Wetzlar, Germany) with an HCX PL APO CS 20X/0.70 dry objective (for YFP in epidermal strips). YFP was excited with the 514 nm laser line, and the 520-560 nm filter was used for emission. Images were later analyzed using ImageJ software (http://rsb.info.nih.gov/ij/) fit-ellipse tool.

### Thermal imaging

Thermal images obtained using an A655SC, FOV 15 (FLIR Systems, Wilsonville, USA). The camera was mounted vertically at about 50 cm above the plants. Images were analyzed by the manufacturer’s software according to the instructions.

### GUS staining

Histochemical detection of GUS activity in was performed using 5-bromo-4-chloro-3-indolyl-β-D-glucuronide as described before (Donnelly et al., 1999). Samples were put on glass cover-slips and photographed under a model ICC50 W bright-field inverted microscope (Leica Microsystem). Images were later analyzed using the ImageJ software. A microscopic ruler (Olympus) was used for size calibration.

### Transport assays

Yeast two-hybrid assays were performed using ProQuest™ Two-Hybrid System with Gateway™ Technology (Thermo Fisher, Waltham, MA, USA). Arabidopsis *ABI1* and *PYR1* cDNAs cloned in pENTR-D-TOPO were introduced into pDEST22 and pDEST32, respectively, by LR reactions. Tomato *AIT1.1* cloned in pENTR-D-TOPO was introduced into pYES-DEST52 of which GAL1 promoter had been replaced with *AHD* promoter (Kanno et al., 2012). Arabidopsis *AIT1* was cloned in pYES-DEST52 in a previous study (Kanno et al., 2012). The pDEST22, pDES32 and pYES-DEST52 constructs were transformed into a yeast strain BY20249. Several (about 10) independent colonies were mixt and precultured in media (SD, -Lus, -Trp, -Ura) overnight at 30 °C and then inoculated on selection media (SD, -Lue, -Trp, -Ura, -His, 1 mM 3AT) with or without (+)-ABA.

For direct measurements of transport activities by LC-MS/MS, AIT1.1 was cloned into the standard pYES-DEST52 vector and transformed into a yeast strain INVSc1. Assays were performed as described previously (Kanno et al., 2016). Hormones were extracted from yeast cells with 1 ml acetone containing 1 % (v/v) acetic acid, and the supernatants after centrifugation were dried up with N_2_ gas. The extracts were dissolved in 1 ml water containing 1 % (v/v) acetic acid and loaded onto cartridge column (Oasis Wax, 1cc; Waters) which had been pre-treated with 0.5 ml acetonitrile and then with water containing 1 % (v/v) acetic acid. After wash with 1 ml water containing 1 % (v/v) and then with 80 % (v/v) acetonitrile, ABA, GA_1_, GA_4_, IAA, Jasmonic acid, jasmonoyl-isolecine were eluted with 80 % (v/v) acetonitrile containing 1 % (v/v) acetic acid. Salicylic acid is then eluted with 2 ml acetonitrile containing 5 % (v/v) formic acid. The eluents containing hormones were dried up with N_2_ gas and the dissolved in 50 µl water containing 1 % (v/v) acetic acid to be analyzed by LC-MS/MS. LC-MS/MS analysis was performed as described previously (Kanno et al., 2016).

### Stomatal aperture measurements

Stomatal aperture was determined using the rapid imprinting technique described by Geisler *et al*. (2000). Light-bodied vinylpolysiloxane dental resin (eliteHD+, Zhermack Clinical, Badia Polesine, Italy) was attached to the abaxial side of the leaflet and then removed as soon as it dried (minutes). The resin epidermal imprints were covered with transparent nail polish, which was removed once it dried and served as a mirror image of the resin imprint. The nail-polish imprints were put on glass cover-slips and photographed under light microscope as described above.

### Stomatal response to ABA

Abaxial epidermal strips were peeled the detached layer were incubated in stomatal opening buffer containing (Wigoda et al., 2006) for 2 h in light conditions (400 µmol m^-2^ sec^-1^). The strips were then transferred to a fresh stomatal opening buffer with 10 µM ABA, in light conditions. After 60 min, the strips were put on glass cover-slips and photographed under the bright-field inverted microscope as described above. Stomatal images were later analyzed using ImageJ for stomatal aperture measurements as described above.

### Statistical analyses

All assays were conducted with three or more biological replicates and analyzed using JMP software (SAS Institute, Cary, NC). Mean comparison was conducted using Tukey-Kramer HSD and Student’s t tests (P<0.05).

### Accession Numbers

Sequence data from this article can be found in the Sol Genomics Network (https://solgenomics.net/) under the following accession numbers: *ACTIN*, Solyc11g005330; *RAB18*, Solyc02g084850; *AIT1.1*, Solyc05g006990; *AIT1.2*, Solyc05g007000; *AIT1.3*, Solyc04g005790; *AIT1.4*, Solyc03g113250.

## Supplemental Data

The following supplemental data are available.

**Supplemental Figure S1.** Molecular Phylogenetic analysis of AIT1 protein sequences in tomato and Arabidopsis.

**Supplemental Figure S2**. PRO activity has no effect on the expression of the tomato *ABCG40* homolog.

**Supplemental Figure S3.** *AIT1.1* promoter drives expression in the vascular tissue.

**Supplemental Figure S4.** Sequence analyses of *ait1.1#1 and ait1.1#7* CRISPR mutants alleles.

**Supplemental Figure S5.** The loss of *AIT1.1* suppressed growth and increased stomatal aperture.

**Supplemental Figure S6.** Plant and leaf phenotypes of 35S:*rgaΔ17 ait1.1*.

**Supplemental Table S1.** List of ABA- and guard cells-related PRO-regulated genes.

**Supplemental Table S2.** gRNAs used in this study.

**Supplemental Table S3**. Primers used in this study.

**Supplemental Dataset S1.** Results of the RNAseq analysis.

## Acknowledgments

We thank Professor Naomi Ori and Professor Yuval Eshed for their valuable discussions and suggestions. We thank Dr. Assaf Mosquna for providing the *MAPKKK18* promoter.

